# Highly versatile small virus-encoded proteins in cellular membranes: A structural perspective on how proteins’ inherent conformational plasticity couples with host membranes’ properties to control cellular processes

**DOI:** 10.1101/2024.08.31.607672

**Authors:** Arvin Saffarian Delkhosh, Elaheh Hadadianpour, Md Majharul Islam, Elka R. Georgieva

**Affiliations:** Department of Chemistry and Biochemistry, Texas Tech University, Lubbock, TX 79409

**Keywords:** viral membrane proteins, viroporins, protein structure, protein conformational dynamics, viral protein-induced membrane permeability to ions

## Abstract

We investigated several small viral proteins that reside and function in cellular membranes, which belong to the viroporin family because they assemble into ion-conducting oligomers. However, despite forming similar oligomeric structures with analogous functions, these proteins have diverse amino acid sequences. In particular, the amino acid compositions of the proposed channel-forming transmembrane (TM) helices are vastly different—some contain residues (e.g., His, Trp, Asp, Ser) that could facilitate cation transport. Still, other voroporins’ TM helices encompass exclusively hydrophobic residues; therefore, it is difficult to explain their channels’ activity, unless other mechanisms (e.g., involving a negative lipid headgroup) take place. For this study, we selected the M2, Vpu, E, p13II, p7, and 2B proteins from the influenza A, HIV-1, human T-cell leukemia, hepatitis C, and picorna viruses, respectively. We discuss the current knowledge of these proteins’ structures as well as remaining questions about a more comprehensive understanding of their structures, conformational dynamics, and function. Finally, we outline strategies to utilize a multi-prong structural approach to overcome current deficiencies in the knowledge about these proteins.

**Highlights:**

- Small viral proteins encoded homo-oligomerize and function in cellular membranes as ion channels
- These proteins were combined in the family of viroporins
- Despite the similarity in their oligomeric structures and functions, these proteins have vastly different primary structures
- It is imperative to understand how proteins with no homology in their primary structures fulfill similar functions for diverse viruses
- There is a need for a multi-prong structural approach to explain the structure, conformational dynamics, and function of these proteins

## 1. Introduction

Pathogenic viruses are among the key environmental factors with a profound negative effect on human health and the sustainable development of human society (https://www.un.org/sustainabledevelopment/). Understanding their mechanisms in hosts is critical for the prevention and overcoming of viral infections. To this end, it is imperative to understand the structure–function relationship of key viral proteins, which will be informative for the design of effective anti-viral therapeutics.

The focus here is on small viral proteins that reside and function in the cellular membranes of infected cells (Figure 1 A). They consist of 50–120 amino acid residues with one or two transmembrane (TM) helices. These proteins are collectively called viroporins because of their ability to form oligomers with ion-conducting channel or pore activities; they also play a role in virus budding.(Seelamgari, Maddukuri et al. 2004, Malim and Emerman 2008, Fischer and Hsu 2011, DiMaio 2014, Luis Nieva and Carrasco 2015, Scott and Griffin 2015, Manzoor, Igarashi et al. 2017, Georgieva 2018, Raheja, George et al. 2023) The influenza A M2 (IAM2) protein was the first viroporin identified and characterized.(Lamb, Zebedee et al. 1985, Sugrue and Hay 1991, Pielak and Chou 2011) After that, viroporins’ family expanded significantly, and such tiny proteins encoded by most currently known viruses have been identified. Besides IAM2, prominent examples of viroporins include the influenza B M2 (IBM2),(Imai, Watanabe et al. 2004) HIV-1 Vpu,(Strebel, Klimkait et al. 1988, Gonzalez 2015) coronavirus (CoV) E,(McClenaghan, Hanson et al. 2020, Santos-Mendoza 2023) hepatitis C virus (HCV) p7,(Chandler, Penin et al. 2012, Atoom, Taylor et al. 2014) and picornavirus 2B (P2B) proteins,(de Jong, de Mattia et al. 2008) as well as human T-cell leukemia virus type I (HTLV-1) p13II protein.(Silic-Benussi, Biasiotto et al. 2010, Georgieva 2018) Commonly, viroporins are subdivided into two classes (Figure 1 B). The Class I consists of members with one TM helix (e.g., IAM2, IBM2, HIV-1 Vpu, CoV E); viroporins with two membrane-traversing helices (e.g., HCV p7, P2B) belong to Class II.(Nieva, Madan et al. 2012) The TM helices of several monomers (the number of monomers is protein-specific) interact in the membrane to form homo-oligomeric structures, which increase membrane permeability for ions.

**Figure 1.**
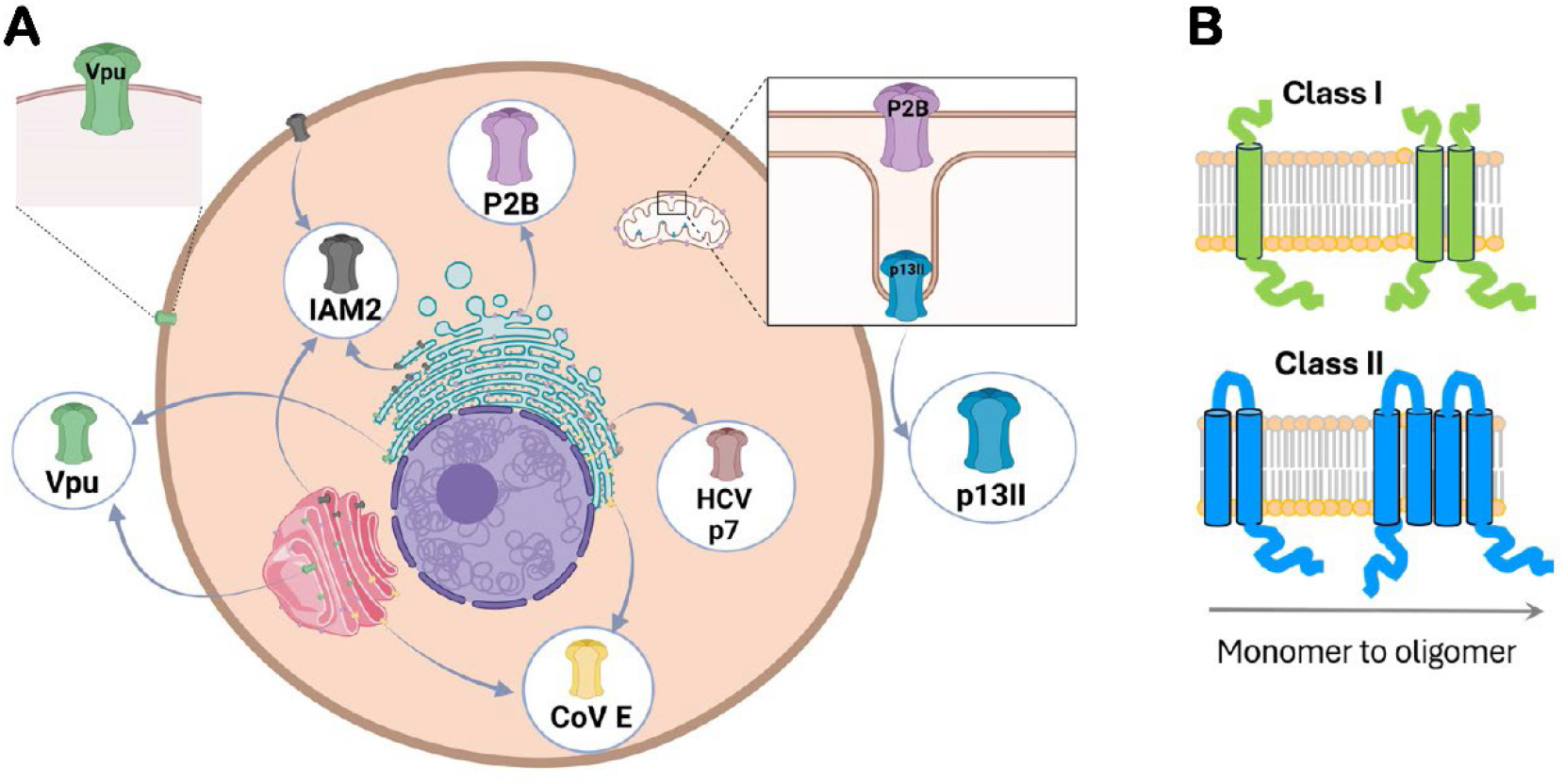
Small viral proteins (viroporins) in host’s cell membranes. (A) These proteins reside and function in plasma and organelle membranes (e.g., membranes of ER and Golgi, and mitochondrial membranes). Furthermore, some of them are found in more than one membrane, e.g., IAM2 and Vpu are located in both plasma and endomembranes (ER and Golgi); P2B is found in endomembranes and outer mitochondrial membrane (OMM). The figure was created in BioRender. (B) The Classes I (1-TM helix) and II (2-TM helices) viroporins are shown; homo-oligomers are formed via TM helix-TM helix associations (only two protomers are shown for clarity).

In addition to their capacity to increase membrane permeability, these tiny proteins participate in diverse protein–protein interactions, forming hetero-protein complexes via their TM helices or soluble domains. Through these interactions, they redirect host signaling pathways by antagonizing host proteins and aid virus budding.(Seelamgari, Maddukuri et al. 2004, Malim and Emerman 2008, Zhu, Wang et al. 2016, Georgieva 2018, Raheja, George et al. 2023) Therefore, these proteins are highly functionally versatile and useful for the respective viruses.

## 2. A brief description of the current status of viroporins’ structural elucidations

The structures of several viroporins have been characterized to various extents. Here, we focus on proteins encoded by six diverse representative viruses—IAM2, HIV-1 Vpu, CoV E, HCV p7, P2B, and HTLV-1 p13II. The amino acid sequences of these proteins and their TM helices are shown in Figure 2. The experimentally determined structures for some of these proteins and/or those generated in this study structures’ predictions using the AlphaFold multimer program(Bryant, Pozzati et al. 2022, Wallner 2023) are shown in Figure 3.

**Figure 2.**
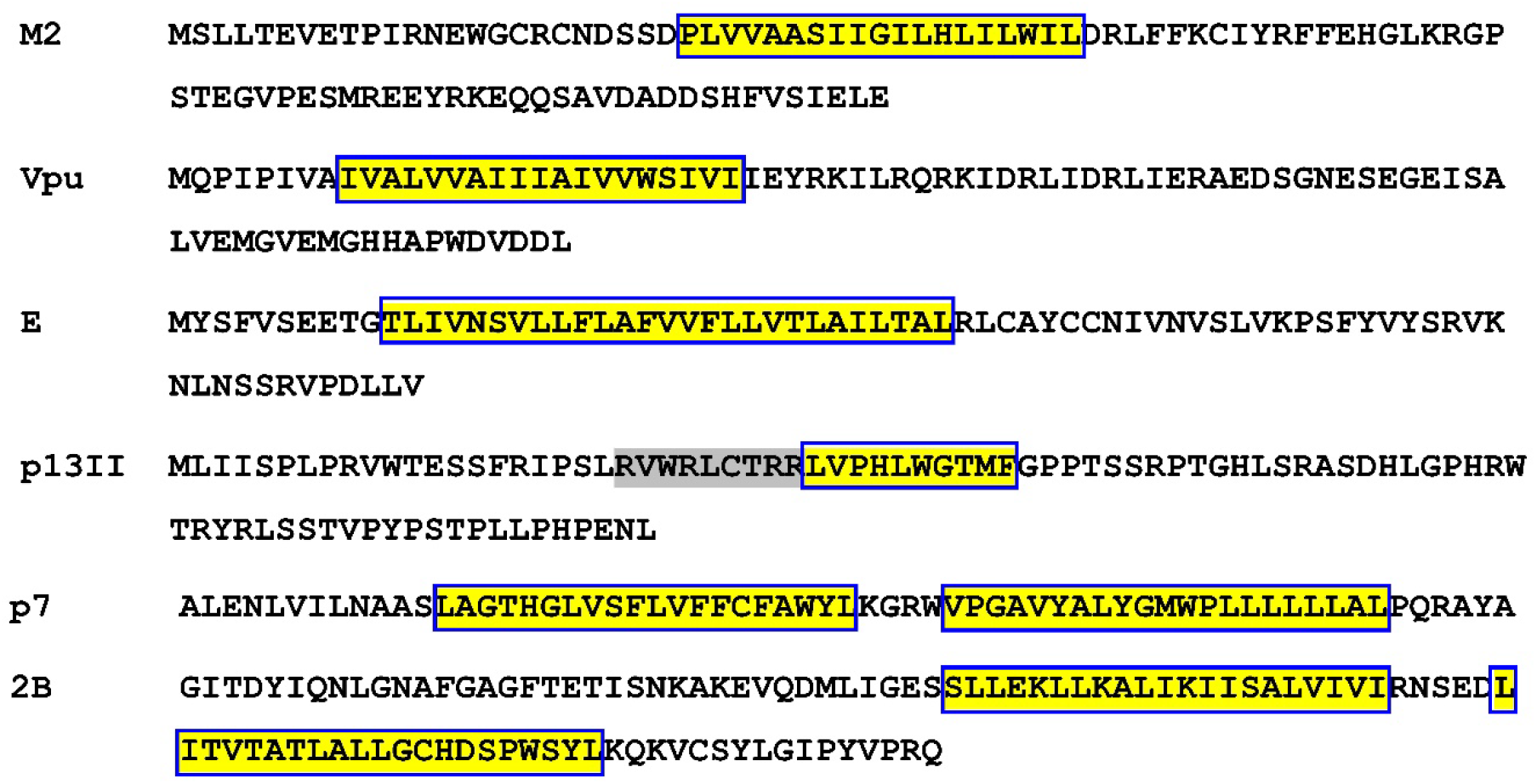
The amino acid sequences of the IAM2 (accession# P63231.1), HIV-1 Vpu (accession#NP_057855.1), CoV E (accession#YP_009724392.1), HTCLV-1 p13II (accession#AAB23362.1), HCV p7 (accession# YP-009709864.1, and P2B (accession# YP-009508946.1) are shown. The TM helices are highlighted in yellow and boxed. The region with positively charged residues in p13II’s sequence is highlighted in gray. The proteins were randomly selected aiming to demonstrate the distinct amino acid composition, particularly the composition of the TM helices, which associate to form ion-conducting channels or pores.

**Figure 3.**
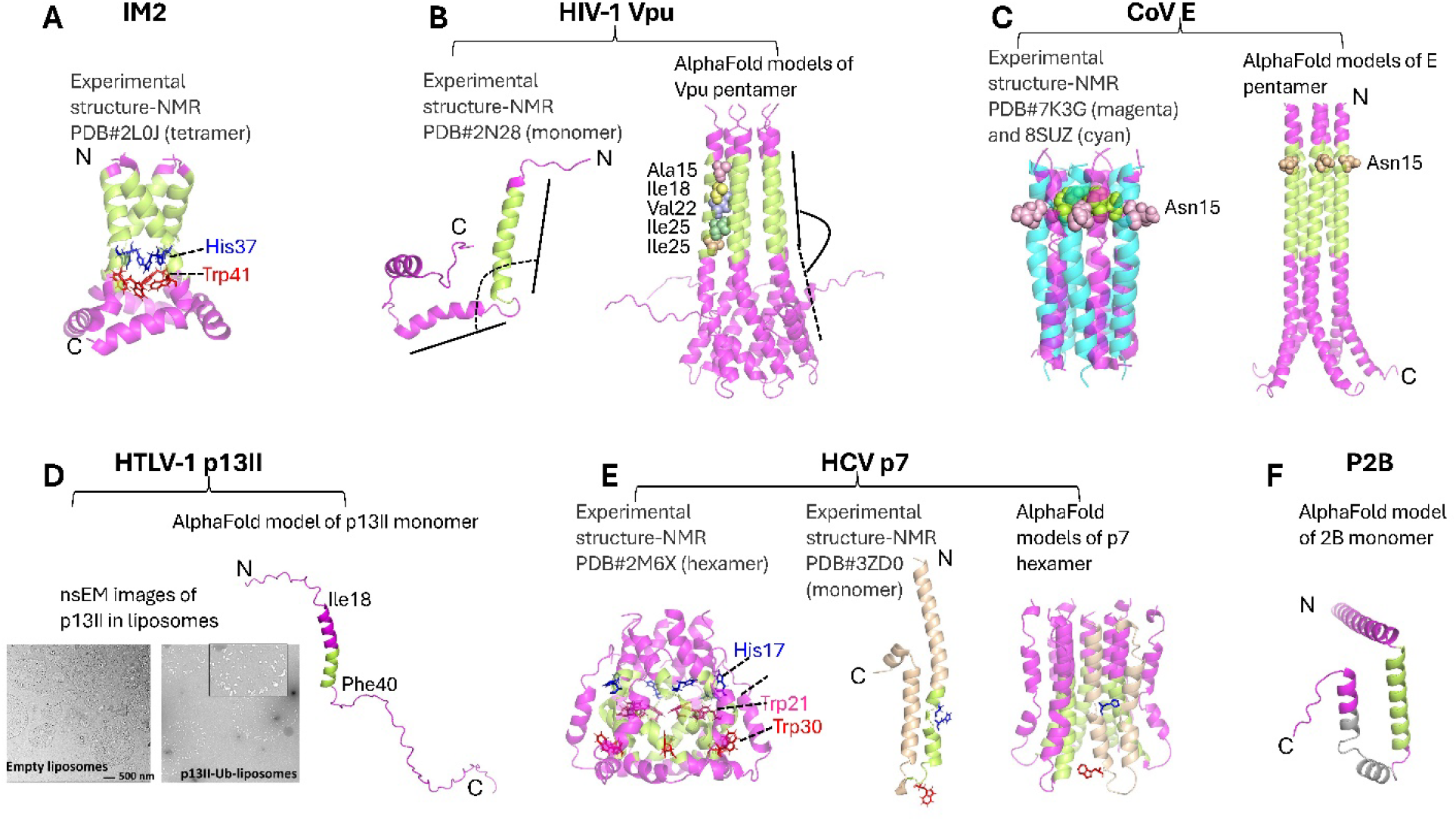
The experimental and predicted structures of IAM2, HIV1 Vpu, CoV E, HICV p7, P2B and HTLV1 p13II proteins. (A) The NMR structure of IAM2 tetramer is shown. The TM helices are in green. The His37 and Thr41 critical for proton’s translocation are shown as sticks in blue and red, respectively. (B) The NMR structure of Vpu monomer (left) and AlphaFold-predicted structure of Vpu pentamer (right) are shown. The TM helices are in green. Based on the NMR structure, helix 2 (after TM helix) is bound to the membrane surface making almost 90° angle with TM helix 1; whereas, based on the AlphaFold prediction, helix 2 is almost a straight continuation of TM helix 1 outside of the membrane. These might correspond to different Vpu conformations, but further clarification would be necessary. Strikingly, all residues in TM helix are highly hydrophobic and how the channel/pore might transport ions is currently unclear. (C) The NMR structures of pentameric E protein TM domain in closed and open conformations are overlayed (left). The experimental results suggest that local restructuring, including helix rotation, at and around residue Asn15 leads to channel/pore opening to facilitate ion translocation. The AlphaFold predicted structure of E protein pentamer is shown. In this model, even in closed state, the Asn15 residue does not point to the inward of the pore. (D) nsEM data of p13II protein reconstituted in DOPC/DOPS liposomes at 1:300 protein-to-lipid molar ratio is shown (left)—The protein reconstitution and nsEM were conducted as described in Georgieva *at al, Prot. Expr. Purif*., 2020 and Majeed *et al, J. Struct. Biol*, 2023, respectively. Large protein clusters were observed. The AlphaFold model of p13II monomer is shown. The predicted TM helix is in green and the amphipathic helix containing several Arg residues is in magenta. (E) The NMR structure of p7 hexamer is shown (left)—unusual fold was observed, the channel is formed with participation of TM helices of each protomer. Another NMR structure of p7 monomer is shown, which is substantially different from the monomer structure in the assembled hexamer (left). This could be because these are the structures of p7 from different genotypes. The AlphaFold model of p7 hexamer is shown on the right. The monomer structure in this oligomer if closer to the structure in the middle. Two conserved residues His17 and Trp30 were identified as possibly participating in ion translocation, but further studies to confirm this are necessary. (F) The AlphaFold model of 2B protein is shown. The predicted TM helices are in green and gray, respectively. All AlphaFold structure predictions were run from ChimeraX.

IAM2 has been a paradigm in studies of channel-/pore-forming homo-oligomeric viral proteins. The protein assembles via its TM helix in a proton-specific tetrameric channel.(Pielak and Chou 2011, Manzoor, Igarashi et al. 2017) A large number tetrameric structures of full-length (FL) of IAM2 and its truncated constructs containing the TM helix were reported by NMR and X-ray crystallography under various lipid, detergent, and pH conditions and supported by MD simulations.(Zhong, Husslein et al. 1998, Duong-Ly, Nanda et al. 2005, Cady, Goodman et al. 2007, Hu, Asbury et al. 2007, Schnell and Chou 2008) In recent studies, Georgieva and colleagues used pulse ESR (DEER) spectroscopy on IAM2 in model liposomes, finding that the tetramer assembles via a tandem mechanism, which is a monomer-to-a dimer-to-a dimer of dimers (tetramer).(Georgieva, Borbat et al. 2015) This mechanism was confirmed for IAM2 reconstituted in isolated *E. coli* membranes containing native lipids and proteins under very close-to-native membrane conditions.(Sanders, Borbat et al. 2024) Additionally, ESR studies shed light on the dynamics of the membrane-bound helix 2 (the IM2 region following the TM helix) of IM2.(Huang, Green et al. 2015)

NMR spectroscopy and X-ray crystallography provided insights into the structure of HIV-1 Vpu protein’s isolated C-terminal region,(Lu, Sharpe et al. 2010, Jia, Weber et al. 2014) TM helix,(Lu, Sharpe et al. 2010) and FL Vpu monomer.(Ma, Marassi et al. 2002) Based on these studies, a pentameric Vpu’s organization in the membrane was proposed and confirmed by computational studies.(Grice, Kerr et al. 1997) Recently, we found that Vpu can self-associate outside of lipid environments, forming predominantly stable hexamers.(Majeed, Adetuyi et al. 2023, Majeed, Dang et al. 2023) However, further investigations would be needed to understand whether Vpu’s soluble and membrane-bound oligomers share similarity.

In lipids, the CoV E protein oligomerizes through its TM helix, possibly as a homopentamer exhibiting ion-channel activity.(Torres, Maheswari et al. 2007, Nieto-Torres, DeDiego et al. 2014, Surya, Li et al. 2018, Schoeman and Fielding 2019, Mandala, McKay et al. 2020) Further computational modeling predicted that the hydrophobic domain (helix) of CoV E could form stable dimers, trimers, and pentamers.(Torres, Wang et al. 2005) An experimentally observed dimeric structure of membrane-bound E protein was reported as well.(Zhang, Qin et al. 2023) Interestingly, a recent study found that in lipid bilayers, the protein can form clusters of pentamers,(Somberg, Wu et al. 2022) similar to the observed IAM2 clusters.(Sutherland, Tran et al. 2022)

The HCV p7 protein localizes primarily on the ER’s membrane with both termini, N- and C-, facing toward the ER lumen,(Khaliq, Jahan et al. 2011) but it also targets mitochondrial membrane.(You, Lee et al. 2017) It belongs to the Class 2 viroporins, as it has two TM helices connected by a short cytoplasmic loop.(Atoom, Taylor et al. 2014) In membranes, p7 forms cation-conducting oligomers (channel or pore).(Chandler, Penin et al. 2012, Atoom, Taylor et al. 2014, You, Lee et al. 2017) It is known from NMR and single-particle electron microscopy (EM) that p7 forms a hexameric unit, with each protomer contributing to ion translocation.(Luik, Chew et al. 2009, Madan and Bartenschlager 2015) However, heptameric(Clarke, Griffin et al. 2006) and pentameric(Pavlovic, Neville et al. 2003) forms of this protein were also reported. Therefore, p7’s oligomerization profile is not well understood, similarly to Vpu and E proteins.

Despite these advances in elucidating the structures and conformational plasticity of several members of the viroporin family, the structures of many of these proteins have not been determined. One such protein is P2B, which has an important role in the life cycle of picornaviruses.(Li, Zou et al. 2019) The structural studies of this protein have not been adequately advanced. It is known from amino acid sequence analysis that this protein has two putative TM helices, TM1 and TM2, arranged in α-helix-turn-α-helix motif.(Li, Zou et al. 2019) It is proposed that the protein assembles in a tetramer,(Agirre, Barco et al. 2002, Patargias, Barke et al. 2009, Li, Zou et al. 2019) the structure of which is currently uncharacterized.

Another interesting protein classified as a viroporin is the HTLV-1 encoded p13II protein,(Silic-Benussi, Biasiotto et al. 2010, Scott and Griffin 2015) which associates with the inner mitochondrial membrane (IMM), where it forms a cation-conducting oligomeric channel.(Silic-Benussi, Cannizzaro et al. 2009, Biasiotto, Aguiari et al. 2010, Silic-Benussi, Biasiotto et al. 2010, Silic-Benussi, Marin et al. 2010) However, very little is known about the structure of this protein in the IMM. The formation of oligomers capable of depolarizing IMM has been reported.(Silic-Benussi, Biasiotto et al. 2010, Silic-Benussi, Marin et al. 2010) For the first time, Georgieva and colleagues reported the production of large quantities highly-pure p13II and found using ESR spectroscopy that this protein oligomerized in lyso lipid and lipid bilayers; the protein also induced Tl^+^ permeability of liposomes, as inferred by fluorescence spectroscopy.(Georgieva, Borbat et al. 2020) Recently, we used negative staining EM (nsEM) to observe large liposome-bound p13II clusters (Figure 3D), which might indicate that membrane permeability is achieved via a different mechanism than those of a channel or pore. Further studies would be needed to characterize these p13II clusters, including solving the structures of the p13II monomer and oligomer.

Thus, in spite of their critical roles in virus adaptation and proliferation, studies on the structure and conformational dynamics of these small membrane-residing viral proteins are incomplete.

## 3. Persisting questions in studies of viroporins

It is currently unknown how viroporins respond structurally to interactions with cellular components (e.g., cellular membranes and host or other viral proteins). It is plausible that these proteins possess high conformational plasticity and can adopt multiple tertiary and quaternary structures to fulfill their diverse functions. However, the small size of these proteins and their membrane localization makes it challenging to assess and characterize each of these structures and to capture the conformational dynamics underlying the transitions between proteins’ functional states. It would be indeed captivating to characterize in detail the varying conformations of these proteins and identify the factors controlling the isomerization between them. In particular, we are interested to understand how viroporins assemble into oligomers to facilitate ion transport across the membrane, and what is the nature of these oligomers. One staggering question is whether a membrane’s permeability increases as a result of the activity of one particular protein’s oligomeric state and whether large proteins’ clusters, such as those observed for IAM2,(Sutherland, Tran et al. 2022) CoV E,(Somberg, Wu et al. 2022) and HTLV1 p13II (Figure 2D), are needed for membrane destabilization. Additionally, there is no clear understanding as to whether these viroporins can form complexes with other protein partners (e.g., host or viral proteins), preserving their channel/pore conformation or the homo-oligomers (and monomers) need to restructure.

Hence, key aspects of the structure–function relationship of these proteins require further elucidation.

Below, we summarize our view on three major persisting questions about channel/pore-forming small viral proteins.

### 3.1. How do proteins with no homology in their amino acid sequences form similar homo-oligomeric structures in the host cellular membranes to fulfil analogous functions?

Careful examination of the primary structures of the proteins from the viroporin family shows no homology among them (Figure 2). Therefore, it is not clear whether these proteins form a family at all, as their amino acid sequences have a little in common, and no evolutionary link is known. It may well be that to gain efficiency, the viruses evolved independently from each other to encode small membrane-residing proteins with similar properties (at least in their capacity to form oligomers in the membrane, thus increasing membrane permeability for ions). A possible approach to put together this fragment of the viroporins’ puzzle is to conduct parallel structural/conformational dynamics studies on several viroporins. This might help to juxtapose the oligomerization mechanisms of a set of proteins, thus uncovering the similarities in their behavior but also singling out the individual properties of each protein.

We believe that oligomerization is an intrinsic property of these small viral membrane proteins, having been observed experimentally for many of the family members. Still, further studies would be needed to link their particular oligomeric structures to specific functions.

### 3.2. How do the amino acid sequences of these proteins couple with the membrane properties to assemble into distinct structures fulfilling specific functions?

Strikingly, unlike the host DNA-encoded TM protein, which resides and functions in specific membranes, each of the functional viroporins is typically found in different environments (e.g., the plasma membrane, ER and Golgi membranes, and OMM and IMM).(Hussain, Das et al. 2007, Silic-Benussi, Biasiotto et al. 2010, Nieto-Torres, Dediego et al. 2011, Gonzalez 2015, Manzoor, Igarashi et al. 2017) This suggests proteins’ high adaptability, as they can adjust to variety of lipid environments (lipid composition, membrane thickness, charge, cholesterol content, etc.), seemingly an advantageous property most likely predefined by their primary structures. Moreover, some of these proteins were found in oligomeric and monomeric forms in different organelle membranes—HIV-1 Vpu protein forms a homopentamer in the membranes of Golgi and intracellular vesicles, but not in the ER’s membrane,(Hussain, Das et al. 2007) confirming that the functional Vpu can transition between multiple quaternary structures.

It is now well understood that the functional oligomeric order of TM proteins is controlled by the helix–helix and helix–lipid interactions (driven by Van der Waals forces, hydrogen bonding, and electrostatic and aromatic interactions).(Cymer, Veerappan et al. 2012, Li, Wimley et al. 2012, Stangl and Schneider 2015) The 3D structure of these proteins is largely determined by their amino acid sequences, but the membrane hydrophobic thickness, fluidity, and charge, which are results of the finely-tuned lipid makeup, are essential for proper folding and assembly of these proteins.(White and Wimley 1999, Corin and Bowie 2022, Levental and Lyman 2023) For viroporins in particular, IM2 tetramer assembles more efficiently in lipids with longer hydrocarbon chains (DOPC/POPS) vs. lipids with shorter hydrocarbon chains (DLPC/DLPS).(Georgieva, Borbat et al. 2016) The IM2’s structural dependence on the length of the lipid hydrocarbon chain was confirmed for protein residing in DMPC vs. DOPC bilayers.(Kovacs, Denny et al. 2000)

Additional studies by others and in our lab show that under the same conditions, the homo-oligomerization properties of various viroporins (e.g., HIV-1Vpu vs. CoV E)(Majeed, Dang et al. 2023, Townsend, Fapohunda et al. 2024) are different. This suggests that each of these proteins uses a subtle individual style to couple its amino acid sequence with the lipid environment to fold into a certain quaternary structure (this possibly affects the tertiary structure as well).

Therefore, from the currently available information, one could assume that the ion-conducting oligomers are formed under particular membrane conditions, but viroporins’ communication with the other viral or host components (e.g., protein-protein hetero-complexes formation) in the infected cell might also initiate the oligomer formation or dissociation.

### 3.3. How can the existing experimental and/or predicted structure of viroporins’ oligomers explain the function of these proteins as ion channels or pores?

Several biophysical and structural biology techniques were utilized to elucidate viroporins’ structures. NMR, X-ray crystallography, and nsEM visualized the IM2 tetramer, Vpu pentamer, E pentamer and p7 hexamer.(Zhong, Husslein et al. 1998, Ma, Marassi et al. 2002, Duong-Ly, Nanda et al. 2005, Schnell and Chou 2008, Luik, Chew et al. 2009, Georgieva, Borbat et al. 2015, Madan and Bartenschlager 2015, Mandala, McKay et al. 2020) Particularly important for understanding the mechanism of ion conductance are the TM domain structures forming the channel/pore across the membrane. It is now well understood that IAM2 is almost exclusively a proton channel, and the conserved His37 and Trp41 (Figure 3A) play critical roles in proton translocation.(Pielak and Chou 2011) A similar arrangement of residue His17 and Trp30 is found in the TM region of the p7 protein (Figure 3E), but no proton specificity or plausible mechanism of ion translocation have been determined for this protein to date. It has been proposed that the E protein channel opens upon an ion entering the pore, triggering TM helix rotation and restructuring in the region of residues N15 and Ser16 (Figure 3C), thus forming a water-filled pore to facilitate the ion translocation. Still, this could explain only halfway of the ion movement in the membrane. The rest of the amino acids in E’s TM are highly hydrophobic and, therefore, may not be suited to provide an ion-translocation pathway. It is even more complex in the case of Vpu—its TM helix’s composition is very hydrophobic (Figure 3B), and there is currently no known mechanism to explain the ion-conducting mechanism of this pentamer. Transient water wires were proposed to mediate ion movement through patches of hydrophobic amino acids in the TM channel.(Kratochvil, Watkins et al. 2023) However, it is not plausible to imagine a water wire spanning 20 hydrophobic amino acids in the case of Vpu. Another proposed mechanism is the formation of a lipid-protein pore, in which the negatively charged lipid headgroups face the pore.(Nieto-Torres, Verdia-Baguena et al. 2015) However, such a structure has not been visualized to date.

Another challenging case is to explain how p13II can span the IMM, given that the proposed TM helix contains only about 10 amino acids (Figure 2 and Figure 3D), which can form a helix with length of about 15 Å, thus insufficient to traverse a membrane with hydrophobic thickness of >25 Å. It is possible that p13II monomers do not fully traverse the membrane but anchor in the bilayer and collectively produce a membrane defect. The region containing several arginines could also play a role in membrane interactions. However, this is yet to be established.

## 4. The need for a multi-prong approach to elucidate the structures and conformational dynamics of proteins from the viroporin family

The importance of the channel-/pore-forming small viral membrane proteins is undisputable because they fulfil key functions in virus adaptation and proliferation in the host. They are also truly captivating proteins because of their functional versatility and conformational adaptability. Still, elucidating these proteins’ structures and conformational dynamics is a challenging task because they are small in size, highly dynamic, and heterogeneous in an oligomeric state (and possibly in tertiary structure too). Last but not least, they reside and function in diverse cellular membranes, imposing additional difficulties to assess their structures. Thus, even in their oligomeric forms, these proteins are out of reach for the single-particle EM analyses and binding of antibodies or protein engineering are required to make these proteins amenable for EM studies.(Nygaard, Kim et al. 2020, Wentinck, Gogou et al. 2022, Majeed, Dang et al. 2023) They are difficult (even impossible) to crystalize or truncated protein constructs and stabilizing mutations need to be introduced, and therefore, X-ray structures might not correspond to the most populated functional protein state.(Thomaston, Alfonso-Prieto et al. 2015, Thomaston, Wu et al. 2019, Sanders, Borbat et al. 2024) NMR can only access certain protein populations because of the requirements for high protein concentration in which an oligomer’s structure is typically modeled;(Ma, Marassi et al. 2002) ESR spectroscopy (CW and pulse [PDS spectroscopy]) can provide information about alteration in local protein dynamics, large-scale structural rearrangements upon transition between conformational states, and the oligomeric order of the protein, but high-resolution structural information is inaccessible with this method.(Jeschke 2012, Georgieva, Borbat et al. 2015, Georgieva 2017, Sahu and Lorigan 2020, Majeed, Ahmad et al. 2021) Molecular dynamics studies(Shukla, Dey et al. 2015, Kolokouris, Kalenderoglou et al. 2021) and protein structure prediction using artificial intelligence-based, (e.g., AlphaFold; Figure 3)(Jumper, Evans et al. 2021) can also help to characterize viroporins’ conformations, oligomeric states, and ion-conduction functions. Still, these proteins have unique amino acid sequences, imposing additional challenges on the homology-based predictions of their structures. Furthermore, AlphaFold has yet to gain the capacity to deal with highly dynamic and heterogeneous proteins in lipid bilayers.(Nussinov, Zhang et al. 2022)

Therefore, it is apparent that the combined effort of several structural and computational biology methods is needed to characterize in detail the multifaced structures of these small viral membrane proteins. In addition, overcoming the current hurdles to study the structures of these proteins will represent a significant methodological contribution in membrane structural biology.

## Author contribution

ASD and EH: literature study, Figure 1, writing the manuscript, data analysis and interpretation; MMI: provided experimental data for p13II (Figure 3 D), data analysis and interpretation; ERG: study conception and design, AlphaFold structures’ modeling, Figures 2 and 3, data analysis and interpretation, writing the manuscript, supervision, funds.

## Acknowledgements

This study was supported by start-up funds (to ERG) and partially by Gilead’s Scholar Program in HIV (to ERG).

“https://www.un.org/sustainabledevelopment/.”

