## Supplementary Information for "Highly versatile small virus-encoded proteins in cellular membranes: A structural perspective on how proteins’ inherent conformational plasticity couples with host membranes’ properties to control cellular processes"

### EXPERIMENTAL PROCEDURES

#### Protein structure modeling

We used the artificial intelligence (AI) AlphaFold2 multimer software (Jumper, Evans et al. 2021) to predict the structures of HIV-1 Vpu pentamer, CoV E pentamer, HTLV-1 p13II monomer, HCV p7 hexamer, and P2B monomer. These structures are shown in Figure 3 in the main text of the manuscript. The AlphaFold program was run in ChimeraX. The structures were visualized using PyMOL software. To the best of our knowledge, these structural models were produces for this particular study and have not been published before.

#### Production, reconstitution in liposomes and nsEM of HTLV-1 p13II protein

The p13II protein was produced as a fusion construct with ubiquitin (p13II-Ub), as previously described (Georgieva, Borbat et al. 2020). The protein was reconstituted in DOPC/POPS (75:25 mol% of non-charged-to-charged lipids) at 1:300 protein-to-lipid molar ratio. The aliquot of 50 mM total lipid stock solution in 20 mM Tris/HCl pH 7.4, 150 mM NaCl, 1 mM EDTA was taken and detergent Triton X-100 (10% stock solution in water) was added until the lipid bilayers were destabilized. The final Triton X-100 concentration was about 1-1.5 %.(Sanders, Borbat et al. 2024) Then calculated volume of p13II-Ub stock solution in a buffer of 20 mM Tris/HCl pH 7.4, 150 mM NaCl, and 100 uM TCEP was added to the destabilized lipid membranes and incubated for 1 h at 22 °C. Thereafter, the detergent was removed using BioBeads, which were changed several times, as previously described (Georgieva, Borbat et al. 2015). After the detergent was removed completely, proteoliposomes and empty liposomes were extruded 15 times using 200 nm polycarbonate membranes and a manual bench-top extruder (Avanti Polar Lipids). In parallel, protein-empty liposomes (with no p13II-Ub) were prepared using the same procedure but just buffer solution was added to the Triton X-100-detabilized lipid instead of adding p13II stock solution.

For negative staining EM (nsEM), we loaded 10µL of the proteoliposome and empty liposome samples onto a carbon-coated copper grid and allowed it to settle for 2 min 20 sec at room temperature. Then, we gently removed the solution on the grid using filter paper and the settled on the grid proteoliposomes and empty liposomes were stained by adding 10 µL of 1.6% 1uranyl acetate (UA) and incubated it for 2 min 20 sec. We again gently cleared the remaining UA solution using filter paper. We air-dried stained samples for 1-2 h at room temperature and then used them for EM imaging. Afterward, we collected digital micrographs with a transmission electron microscope (TEM Hitachi H-7650), as described previously (Majeed, Adetuyi et al. 2023).

The nsEM images of empty liposomes and proteoliposomes are shown in Figure 3 D in the main text.
